# The Primary Tumor Immune Microenviornment Status Predicts the Response to Immunotherapy and Overall Survival in Breast Cancer

**DOI:** 10.1101/2020.07.03.186221

**Authors:** Arjun Moorthy, Aidan Quinn

**Affiliations:** Imaging Endpoints, LLC Scottsdale, AZ; BASIS Scottsdale, Scottsdale, AZ; Department of Pathology and Cell Biology, Columbia University, New York, NY; Institute for Cancer Genetics, Columbia University Medical Center, New York, NY

## Abstract

The tumor immune microenvironment (TIME) of breast cancer is a known source of tumor heterogeneity and it has been increasingly recognized as having a role in the course of disease. In the present study, we used a computational approach to dissect the landscape of TIME states among TCGA breast cancer patients. Our central hypothesis is that the pre-existing TIME states represent a dimension which is informative about the prognosis and the response to immunotherapy. In order to test this hypothesis, we first classified breast cancer patients according to their primary TIME status. Next, we describe a TIME-based classification with prognostic value for overall survival among the TCGA patients. We further demonstrated that absolute quantification of mast cells, M0 macrophages, CD8 T cells and neutrophils were predictive of overall survival. In order to identify the TIME states which, predict response to immune checkpoint blockade, we performed a similar analysis of 11 different mouse models of primary invasive breast carcinoma that were subsequently treated with immune checkpoint inhibitor (ICI) therapy. These analyses revealed that the TIME content of M1 macrophages, monocytes and resting dendritic cells were predictive of sensitivity to ICI therapy. Taken together, these results indicate that (1) the landscape of human primary TIME states is diverse and can identify patients with more or less aggressive disease and (2) that pre-existing TIME states may be able to identify patients, of all molecular subtypes of breast cancer, who are good candidates for ICI therapy.

## III. Introduction

Breast invasive adenocarcinoma is the most frequently diagnosed malignancy among women in the United States, in 2020 it is expected to account for 15.3% of all new cancer diagnoses and 42,170 of all cancer deaths.[1] Due largely to the development of aggressive and improved treatment strategies as well as several targeted therapeutic agents, over the past two decades the death rate of breast cancer has declined from about 31% in 1992 to 20% in 2017 with a 5 year survival rate of 90% between 2010-2016.[1] Despite these recent advances, resistance to all known therapeutics still occurs in some women, especially those with basal subtype tumors defined as HER2, PR and ER negative (or triple negative breast cancer), and these patients inexorably progress in their disease.[2]

Notably, however, immunotherapies such as immune checkpoint inhibition (ICI) in particular, have shown enormous potential in treating otherwise incurable carcinomas including metastatic melanoma and lung cancer.[3,4] Recently, ICI treatments have proven to extend survival among TNBC patients with metastatic disease. The IMpassion130 trial[5], demonstrated that the anti-PDL1 therapy, atezolizumab in combination with nab-paclitaxel extended overall survival compared to nab-paclitaxel alone among patients with tumors that express PDL-1. However, the response rate among even patients with expression of PDL-1 was variable and expression of PDL-1 alone is unlikely to fully account for the full spectrum of responses to atezolizumab. Moreover, the IMpassion130 trial demonstrated that patients with metastatic disease limited to lymph nodes received a far greater benefit from the atezolizumab, nab-paclitaxel combination compared to those with distant metastases indicating a potential role for ICI therapy at earlier stages in the disease. Taken together, these observations suggest that breast cancer patient selection for ICI intervention is of critical importance and should be studied in detail.

ICI therapies block the inactivation of the anti-tumor immune response by the tumor itself, thus promoting immune-mediated cell killing of the tumor. Therefore, the primary tumor immune microenvironment (TIME) may have a role in determining the effect of ICI therapies in breast cancer by establishing a permissive or suppressive microenvironment for the immune system thereby adding to or detracting from the effect of ICI therapy, respectively.

Recent efforts to identify biomarkers for and mechanisms of resistance to ICI therapy have focused primarily on genetic or tumor intrinsic modes including the mutational burdon of the tumor[6] and the expression of ICI target molecules and immune modulating genes including PDL-1 itself [7–9]. Relatively little, however, is known about the exact cellular composition of the primary TIME of breast cancer in general and which TIME states specifically are predictive of ICI response. Early work in identifying the TIME determinants of response to ICI therapy has demonstrated the importance of the tumor lymphocyte (TIL) and macrophage abundance [10–12] However, with recent the development of single cell high-throughput sequencing, studies have begun to dissect the breast cancer TIME at the cell type compositional level [13,14]. Despite this technological advancement, high cost and computational constraints remain and it is largely infeasible to perform these experiments at the scale required to achieve the statistical power to characterize the complete landscape of TIME statuses present among large, heterogenous cohorts of breast cancer patients and associate trends in this landscape with clinical outcomes.

In the present study, we apply CIBERSORT [15], a computational approach to infer the abundance of specific cell types from bulk RNA-seq data, to 922 individual samples of human primary breast cancer from TCGA [16]. Using this approach, we were able to interrogate the trends in the TIME statuses of these patients that are associated with prognosis of the disease. In addition, we were able to apply a similar approach to mouse models of metastatic breast cancer to identify TIME states that are predictive of objective response to ICI therapy in these models.

## IV. Materials and Methods

### Human Primary Tumor RNA-seq Data

Raw RNA-seq data and associated clinical metadata for 1,035 female patients with primary breast invasive carcinoma in the TCGA database was downloaded using the GDC Data Portal. Molecular subtype (Her2, Basal, or Luminal) was determined for each case using estrogen receptor, progesterone receptor and HER2 status. Cases with equivocal or absent histological measurements were excluded from further analysis (n = 113). Gene level summarized RNA-seq read counts were normalized and statistical analyses performed using the R package DESeq2 [17].

### Mouse Primary Tumor RNA-seq Data

Pre-treatment RNA-seq data from 47 individual animals comprising 11 different mouse models of triple negative breast cancer was obtained from GEO (GSE124821). Briefly, and as detailed by Hollern *et al*. in the original work [18], these animals were subsequently treated with anti-CTLA4 and anti-PDL1 antibodies (bi-weekly, intraperitoneal injection) and objective response to this combination therapy was recorded.

### Inference of Tumor Microenvironment Content from Bulk RNA-seq

Scaled and normalized RNA-seq reads were imported into CIBERSORT [15] and TIME cell content was inferred for all populations in the LM22 gene signature database. Immune cell quantitation was performed with default parameters across 500 permutations run in relative and absolute modes. Absolute quantification files were imported into R for downstream statistical analysis and plotting.

### Statistical Analysis

Survival analysis was performed in R using the survival package and Kaplan-Meier plots were generated using the survminer package. All statistical tests were computed using base R v4.0.1.

## V. Results

### TIME Landscape of Human Primary Invasive Breast Carcinoma

In depth analysis and characterization of the TIME landscape of 1,035 primary samples of human breast invasive carcinoma revealed profound variability in the constituent immune cell types among patients (Figure 1a). Notably, high levels of all broadly detectable immune cell types (macrophages, monocytes, resting mast cells, CD4 and CD8 T cells, B cells) were detected in approximately 25% of all primary tumors - a feature indicative of immunologically hot tumors, while 12% had low to undetectable levels (> 2 *s*.*d*. below the population mean) - indicative of immunologically cold tumors. M2 macrophages and CD4 T resting memory cells were the most frequently detected immune cell component of the breast cancer immune microenvironment, detected in 98% and 92% of primary tumors, respectively. As expected, rare immunological cell types such as neutrophils, eosinophils, activated mast cells, naive CD4 T cells, memory B cells and γδ T cells were only detected above background in 0-3% of primary tumors. Taken together, these results indicate that CIBERSORT deconvolution of the TIME of human invasive breast carcinoma recapitulates the expected variation among individuals and distribution of specific cell types.

**Figure 1.**
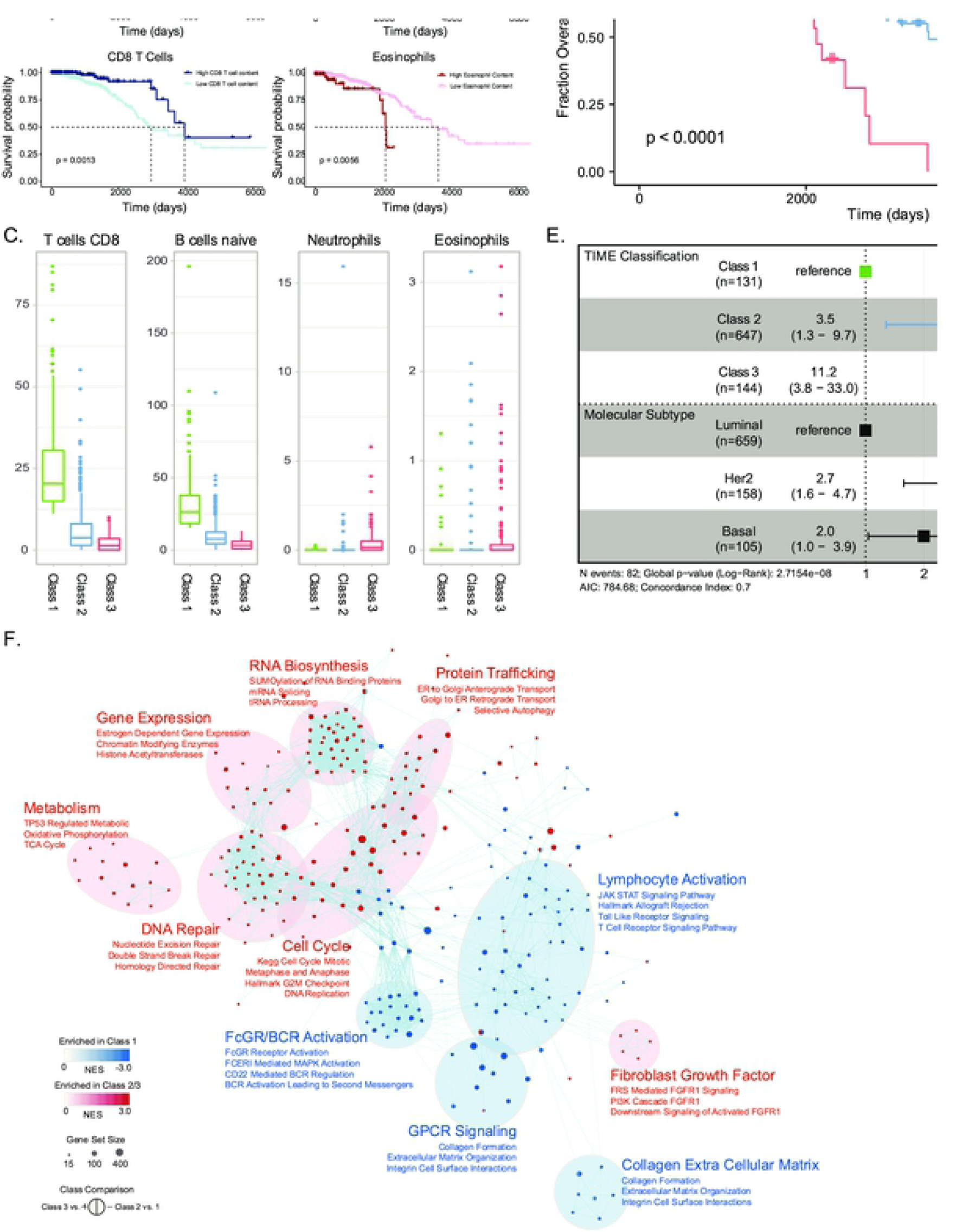
Primary TIME States Segregate Human Invasive Breast Carcinoma into TIME Classes with Functionally and Prognostically Distinct Features. A. Heatmap of the primary TIME landscape of human metastatic breast carcinoma. All 22 identifiable cell types are displayed on the vertical axis and individual patients are clustered along the horizontal axis. Column colors indicated the molecular subtype and the annotated front-line treatment type for each patient. B. Kaplan-Meyer plots for individual TIME cell types most predictive of overall survival. C. Distributions of the absolute qualification of the key populations of TIME cells across the three TIME classes. D. Kaplan-Meyer plot comparing the overall survival among patients in the three TIME classes. E. Cox-PH regression model testing the prognostic value of TIME classification and molecular subtype. TIME-classification was predictive of overall survival independently of molecular subtype. F. Gene set enrichment analysis comparing the transcriptional profiles of aggressive TIME-classes 2 and 3 to class 1.

### TIME Features are Predictive of Prognosis in Primary Human Breast Tumors

Given that the presence and/or absence of immune cells in the TIME have been shown to influence the course of disease including invasiveness, metastasis and prognosis [19,20], we next determined the relationship between overall survival and the TIME content of each individual cell type within the LM22 signature set. The presence of high levels (> 25th percentile) of specific lymphocyte types, Naive B cells (*p*=0.0033) and CD8 T cells (*p*=0.0013), conferred a significantly better prognosis compared to patients with lower TIME content of these cells (Figure 1b). Interestingly, patients with any detectable TIME content of neutrophils (*p*=0.0056) and eosinophils (*p*=0.0056) had significantly worse prognosis compared to those without.

Based on the predictive capacity of lymphocytes and granulocytes in the primary breast TIME, we designed a stronger classifier by aggregating the individual cell content information for each of these individual cell types - which function as weak classifiers in our model. Patients with naïve B cell content and CD8 T cell content higher than the 25th percentile for all patients and undetectable neutrophil or eosinophil content were assigned to class 1 (*n* = 131). Patients with the inverse TIME profiles were assigned to class 3 (*n* = 144), and patients that failed to fit into either of these two groups were assigned to an intermediate class 2 (*n* = 647).

As expected, average CD8 T Cell and naïve B cell content were significantly higher in class 1 patients compared to class 3 (*p* < 2.2e-16, Figure 1c). Class 2 patients tended to have greater numbers of CD8 T cells and naïve B cells compared to those in class 3 (*p* < 2.2e-16), but significantly lower than those in class 1 (*p* < 2.2e-16). Neutrophil and Eosinophil content was low in both class 1 and class 2 patients but only significantly higher in class 3 compared to both classes 1 and 2 (*p* = 8.08e-10 and p = 0.0011, respectively).

Consistent with the hypothesis that primary TIME status of primary tumors is predictive of outcomes, our primary TIME classification strategy was able to identify patients significantly different overall survival (p < 0.0001, Figure 1d). Moreover, this predictive capacity of the primary TIME status remained statistically significant after controlling for molecular subtype (Figure 1e). Patients with class 1 primary tumors had the best prognosis with a median overall survival of 11 years and greater than 90% survival rate beyond 9.6 years. Patients with class 2 tumors had an intermediate prognosis with a median survival of 9.51 years and patients with class 3 tumors had the worst prognosis with a median overall survival of only 5.83 years and fewer than 15% achieving long-term survival beyond 8 years. After controlling for molecular subtype, patients with class 2 tumors had hazard ratio of 3.5 (95% CI: 1.3 - 9.7, *p* = 0.015), compared to those with class 1 tumors, and patients with class 3 tumors had a hazard ratio of 11.2 (95% CI: 3.8 - 33.0, *p* < 0.001).

### Primary TIME Status is Predictive of Response to ICI Therapy

Given the capacity of the human primary TIME to predict prognosis in the context of conventional therapeutic strategies, we next sought to test whether the primary TIME was informative for the response to immune checkpoint inhibition (ICI) therapy (Figure 2a). CIBERSORT analysis was performed on publicly available RNA-seq data obtained from a panel of 11 different mouse models of triple negative breast cancer, which were subsequently treated with a combination of anti-PDL1 and anti-CTLA4 ICI and an objective response was measured. This analysis revealed a heterogeneous primary TIME, qualitatively similar to that of the human breast cancer tumors obtained from TCGA (Figure 2b).

**Figure 2.**
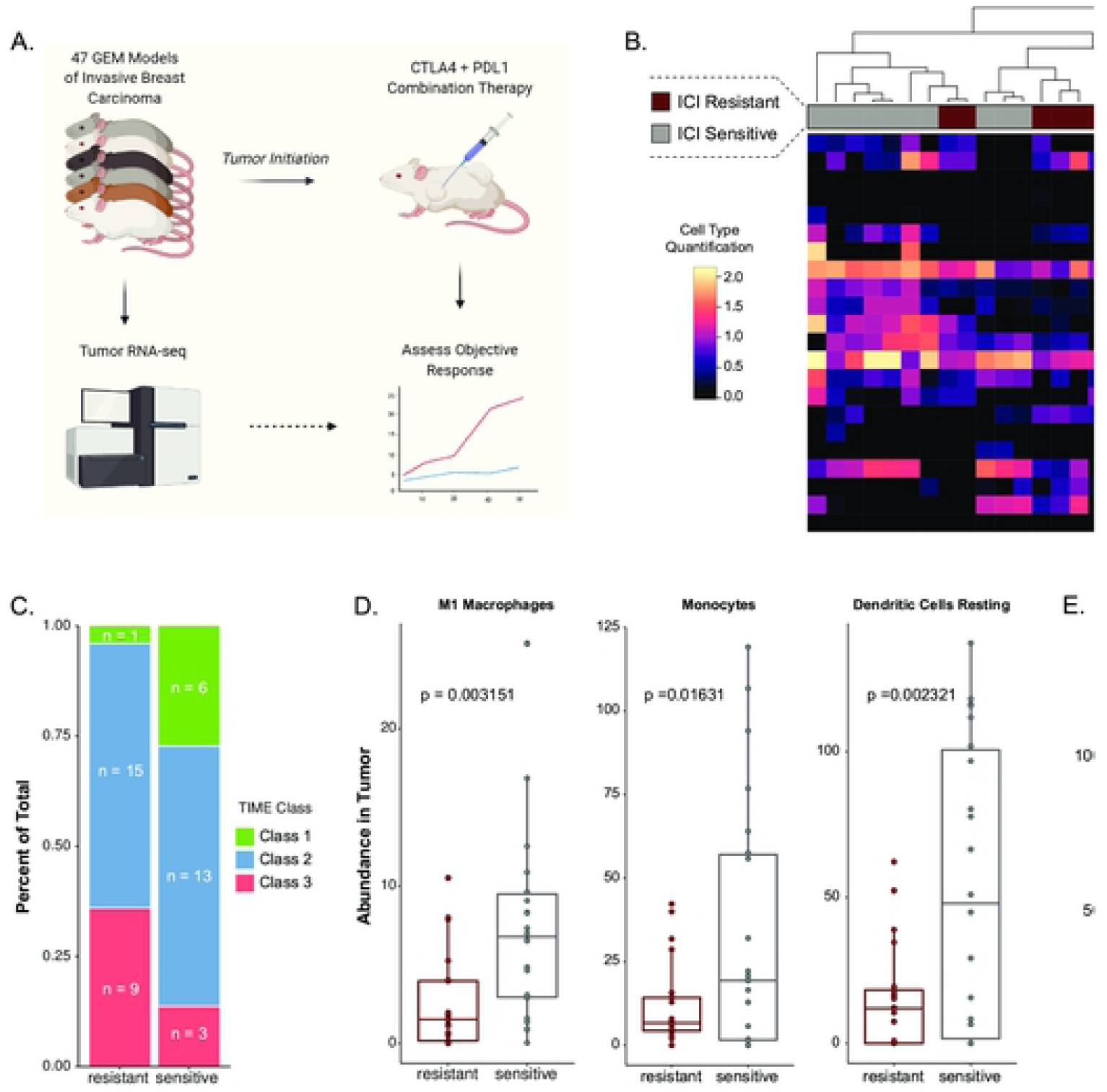
Primary TIME Status Predicts Response to Immune Checkpoint Inhibition Therapy in Mouse Models of Breast Cancer. A. Schematic of the experimental approach used by Hollern *et al*. to generate the database of murine models of breast cancer response to ICI treatment. B. Primary TIME landscape of the mouse models of invasive breast cancer; cell types are displayed on the vertical axis and individual mice are clustered along the horizontal axis. Column colors indicated the response to ICI treatment type for each animal. C. Percent of total mice with primary TIME class (class 1, 2 or 3) separated by response to ICI therapy. D. Distribution of the absolute quantification of the TIME cells most predictive of response among ICI sensitive and resistant mice. E. Distribution of the ICI Response Scores of individual mice separated by ICI response. F. ICI Response score for all 922 primary human tumor samples included in the TCGA analysis separated by TIME class (class 1, 2 and 3).

Notable similarities to the human data include; high TIME content of M0 macrophages was identified among approximately 65% of mouse tumors, M2 macrophage content was consistently high in all tumors, high TIME content of plasma cells were identified in a minority of tumors (<10%), and tumors were largely devoid of rare immune cell populations such as neutrophils.

We next determined the TIME class of the mouse models according to our method described above. The distribution of TIME classifications of these mouse models closely resembled the distribution of time classifications among the human tumors, with class 1 tumors comprising 14.9%, class 2 comprising 59.5% and class 3 comprising 25.5% of mouse tumors. As expected, the TIME class of the ICI-sensitive mice was lower on average compared to those that were resistant to ICI therapy (figure 2c).

Consistent with other reports, our analysis revealed that TIME content of lymphocytes was predictive of the objective response to ICI treatment (supplemental figure 1). Specifically, plasma cell, CD8 T cell and CD4 memory T cell contents were significantly associated with response.

### TIME Based ICI Response Score

In order to more quantitatively and sensitively identify TIME statuses which could be predictive of ICI therapy response, we next compared the predictive capacity of each individual TIME components and developed an ICI Response Score (RS) on the basis of the top individual predictors - M1 macrophages, monocytes and resting dendritic cells (figure 2d).

Briefly, the CIBERSORT absolute quantifications for each sample *i*, were used to compute the quantity

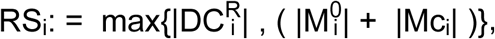

where DC^R^ is the resting dendritic cell content, M^0^ is the M0 macrophage content and Mc is the monocyte content.

The ICI Response Score of the primary TIME was significantly higher among animals that objectively responded (mean RS of 25, +/-13 *s*.*d*.) compared to those that did not respond (mean RS = 75, +/-20 *s*.*d*., *p* = 2.1×10^−6^), indicating that this response score may be useful to identify breast cancer patients who could benefit from ICI therapy.

Next, to determine whether RS was associated with a particular TIME class in human patients, we computed the RS for each patient in our TCGA data set (figure 2f). This analysis revealed that patients with class 3 primary TIME status had significantly lower RS, compared to those in class 1 (*p* < 0.001). However, we were able to identify patients of all primary TIME classes and molecular subtypes with high RS’s indicating that this subset of breast cancer patients might respond well to ICI therapy.

## VI. Discussion

The role of the tumor immune microenvironment in the course of disease of breast cancer including prognosis and response to cancer immunotherapies remains poorly understood. In this study we used an unbiased computational approach to infer the cellular composition of the primary TIME of 922 patients from bulk RNA-seq data. From this rich dataset we were able to identify primary TIME states that are informative for prognosis indecently of molecular subtype in human breast cancer. In addition, we identified pre-existing TIME states that are related to response to immune checkpoint inhibition in a panel of mouse models of invasive breast carcinoma, indicating the utility of this approach to dissecting the primary TIME for predicting response to ICI therapy.

Previous studies aimed at elucidating the primary breast cancer TIME have generally either examined a small subset of TIME components among a large cohort of patients or taken a less biased approach such as single cell RNA-seq to carefully dissect the TIME status of smaller numbers patients [21]. Here, using computational methods to infer the absolute quantifications of 22 different immune cell populations from the bulk RNA-sequencing data we were able to perform a less biased study of a large cohort of patients. This approach allowed us to identify a highly variable landscape of primary TIME states among the TCGA breast cancer patients. Suggesting that there is a high degree of inter-tumoral variability in the TIME content, which has been postulated to underly differential responses to both traditional and immunotherapies [22].

Specifically, the relationship between tumor lymphocyte content and overall survival are supported by our data, however we additionally identify that high neutrophil and eosinophil content confers a worse overall prognosis. Our novel primary TIME-based classification strategy incorporates these findings and demonstrates that the TIME state of the primary tumor indeed has prognostic value, independently of the molecular subtype of the tumor.

Furthermore, in order to use the primary TIME status to identify those patients who could benefit from immune checkpoint inhibition therapy, we developed an ICI Response Scored based on primary TIME status. A subset of patients of all molecular subtypes and primary TIME classes were identified with high ICI Response Scores, suggesting that future ICI therapy preclinical and clinical trials would benefit from stratification methods which consider the primary TIME prior to enrolment.

Taken together, these findings demonstrate that pre-existing TIME states are relevant to both the prognosis of breast cancer patients and to the choice of therapy. Future studies should further dissect these TIME states by flow cytometric and/or single cell RNA-sequencing approaches in prospective studies.

